# ODDPub – a Text-Mining Algorithm to Detect Data Sharing in Biomedical Publications

**DOI:** 10.1101/2020.05.11.088021

**Authors:** Nico Riedel, Miriam Kip, Evgeny Bobrov

## Abstract

Open research data are increasingly recognized as a quality indicator and an important resource to increase transparency, robustness and collaboration in science. However, no standardized way of reporting Open Data in publications exists, making it difficult to find shared datasets and assess the prevalence of Open Data in an automated fashion.

We developed ODDPub (Open Data Detection in Publications), a text-mining algorithm that screens biomedical publications and detects cases of Open Data. Using English-language original research publications from a single biomedical research institution (n=8689) and randomly selected from PubMed (n=1500) we iteratively developed a set of derived keyword categories. ODDPub can detect data sharing through field-specific repositories, general-purpose repositories or the supplement. Additionally, it can detect shared analysis code (Open Code).

To validate ODDPub, we manually screened 792 publications randomly selected from PubMed. On this validation dataset, our algorithm detected Open Data publications with a sensitivity of 0.74 and specificity of 0.97. Open Data was detected for 11.5% (n=91) of publications. Open Code was detected for 1.4% (n=11) of publications with a sensitivity of 0.73 and specificity of 1.00. We compared our results to the linked datasets found in the databases PubMed and Web of Science.

Our algorithm can automatically screen large numbers of publications for Open Data. It can thus be used to assess Open Data sharing rates on the level of subject areas, journals, or institutions. It can also identify individual Open Data publications in a larger publication corpus. ODDPub is published as an R package on GitHub.

## Introduction

The benefits of openly shared research data (Open Data) for science and society as a whole are manifold, including increases in reproducibility, resource efficiency, economic growth, and public trust in research, as stated e.g. by the Concordat on Open Research Data, drafted by Universities UK, Research Councils UK, HEFCE and Wellcome Trust (The Concordat Working Group 2016). Open Data are increasingly recognized as both an indicator of quality and an important resource that can be reused by other scientists and that can increase transparency, robustness, and collaboration in science (Fecher, Friesike, and Hebing 2015). Several large funders and institutions advocate for Open Data, e.g. the European Commission, NIH, EUA, RDA Europe (Guedj and Ramjoué 2015; NIH 2003; EUA 2017). The Berlin Declaration on Open Access to Knowledge has been signed by over 600 institutions and organizations (“The Berlin Declaration on Open Access to Knowledge” 2003) and the Sorbonne declaration on research data rights was signed recently and is very widely supported (“Sorbonne Declaration on Research Data Rights” 2020). Correspondingly, large infrastructure projects are under way, including the European Open Science Cloud (“EOSC Declaration” 2017). Similarly, Open Code, i.e. analysis code disseminated with the publication, is recognized as an element to further enhance transparency of a study by making available the analysis steps leading from data to results, which is increasingly reflected in journal policies (Stodden, Guo, and Ma 2013).

On the level of individual researchers, Open Data can lead to increased visibility, resource efficiency (Pronk 2019; Milham et al. 2018), and possibly a citation advantage (Colavizza et al. 2020; Piwowar and Vision 2013). However, there are also limits to the open sharing of research data in the biomedical domain, especially when it comes to patient data that are subject to data protection regulations. In those cases, only restricted forms of data sharing involving application procedures and/or deidentification might be possible (Tudur Smith et al. 2015).

Academic institutions have a role to play in incentivizing Open Science by assessing research quality and transparency for the allocation of funds, hiring or promotion. The inclusion of Open Science metrics such as the prevalence of Open Data or Open Code in institutional incentive and reward structures is a necessary step to increase Open Science practices (Strech et al. 2020).

Shared research data allows researchers to find and use already existing datasets. There are currently several databases listing datasets linked to publications. PubMed and Web of Science both provide metadata fields that link shared datasets to publications. Additionally, there are search engines that are specifically focused on openly available datasets, like DataCite, Datamed or Google Dataset Search.

However, there is not yet a standardized way of sharing and reporting Open Data, which makes it difficult to find and link these datasets. Data are currently shared in many different ways, in different field-specific or general-purpose repositories or with data shared as the supplement of a publication. Additionally, the reporting of data sharing is very heterogeneous in the literature. While some journals and publishers like PLoS or BMC have adopted policies on data sharing and introduced mandatory data availability statements, many other journals do not request such statement. This splits the literature into journals in which data availability is relatively easy to check and journals where data availability is not reported in a standardized way. Current studies on data sharing mostly focus on specific journals with mandatory data availability statements (Federer et al. 2018; Colavizza et al. 2020). Open Data may also be shared together with the publication, e.g. in the supplement, or as an independent digital object and cited in the publication (Cousijn et al. 2018; Martone 2014).

As currently only a minority of journals have mandatory data availability statements and the sharing of data as independent digital objects is not common, it is difficult to assess the prevalence of Open Data for the broad biomedical literature. If we want to monitor Open Data prevalence on the level of research fields or institutions, an approach that does not only rely on data availability statements but rather searches the whole publication texts is necessary. Monitoring on the institutional level would also be important to assess the success of an institutional Open Data policy or the effectiveness of incentives for Open Data practices.

In this publication, we develop and validate the text-mining algorithm ODDPub (Open Data Detection in Publications), which screens biomedical publications for Open Data statements. With this approach, ODDPub allows to obtain estimates of the prevalence of Open Data in the biomedical literature on a larger scale and independent of data availability statements. The ODDPub algorithm is freely available as R package on GitHub (https://doi.org/10.5281/zenodo.3760970, RRID:SCR_018385).

## Methods

### Definition for Open Data & Open Code used in this study

Apart from openness as defined by the Open Definition, see (Open Knowledge Foundation 2017), there are other criteria concerning the reusability of datasets, known as the FAIR criteria (Wilkinson et al. 2016). However, few publications would currently comply with the full set of FAIR criteria. Therefore, we used a definition of Open Data that covers only some aspects of the FAIR criteria in addition to the openness criterion.

We only considered research data shared together with a publication. These research data could be raw or preprocessed, but not analyzed or aggregated. Thus, they would allow the analytical replication of the results (or at least a part thereof). However, an imprecision with respect to when data are considered “raw” necessarily remains due to differences in disciplinary practices. Primary data or secondary data (e.g. resulting from the combination of freely available data sets or from meta-analysis) were both eligible, as long as they fulfilled above requirement and the datasets shared were generated by the authors. We considered image data or other audiovisual data only if they encompass a substantial amount of the research data used in the study. In case of doubt, data deposited in repositories were considered Open Data due to their findability and potential reusability, while conversely data in supplementary materials were only considered Open Data if these were clearly raw data. We excluded audiovisual data that are primarily shared for illustrative purposes. The amount of data was not a criterion for inclusion per se, such that data from a study with a handful of subjects were still considered Open Data.

We used similar criteria for Open Code, and counted a publication as Open Data/Code publication if:

i. The publication explicitly mentioned where the shared data/code can be found. This can be a DOI, URL, database ID/accession code or a reference to a specific file in the supplement (a reference to supplementary materials without further details was not sufficient)
ii. The data/code was freely accessible to everyone. Data/code was not only shared upon request and no application or registration process was required.
iii. The data/code in principle allowed an analytical replication of at least parts of the results. Sharing summary results was not sufficient. However, we did not check the actual analytical replicability of the publications, as this is outside the scope of this study.
iv. The data/code was in a format that is machine-readable or that can easily be transferred into one. We allowed data shared in Excel- or Word-Tables but excluded data shared in PDFs (including the main publication manuscript) or as pictures of tables. If the data were shared in a discipline-specific format, we considered them reusable.
v. The reuse of data/code previously published by other researchers was not considered.
vi. We did not presuppose an explicitly stated license (e.g. CC BY, MIT) for the shared data/code.

We specifically did not address all possible kinds of data sharing statements. We explicitly excluded cases of data sharing where a registration or application is necessary to access the data. While cases of restricted access to data can have good reasons, for example for clinical datasets, this is out of the scope of the current work, which only considers Open Data. We also excluded cases of data being available upon request. Making data available upon request is considered a suboptimal data sharing strategy and several studies (Savage and Vickers 2009; Rowhani-Farid and Barnett 2016; Naudet et al. 2018) have shown that in many of those cases data cannot be accessed, especially after longer periods of time (Vines et al. 2014).

### Creation of training and validation datasets

We created two training sets and a validation dataset for the development of the algorithm.

The first training dataset consisted of the institutional publications record of the Charité - Universitätsmedizin Berlin from the years 2015 to 2017. To retrieve the publication record, we searched the publication databases PubMed and Embase for publications with affiliations that matched a list of keywords with variations of the institutional name (see Supplementary File S1). Additionally, we filtered the results for the publication years 2015-17 and for the publication type ‘journal article’ (PubMed) or ‘article’ (Embase). We obtained 9896 results from PubMed and 9256 results from Embase. The resulting publication lists from both databases were then combined and duplicates were removed by matching the DOI, PubMed ID (PMID), or the title and the last name of the first author. The combined publication list consisted of 11192 entries. To retrieve the publication PDFs, we loaded the combined publication list into EndNote^™^ (Version: X8.2, https://www.endnote.com) and used the ‘Find full text’ function. That way, 8689 (78%) of the publications could be retrieved. Some full texts could not be retrieved from journals for which our institution has no access (see Figure 1).

**Figure 1:**
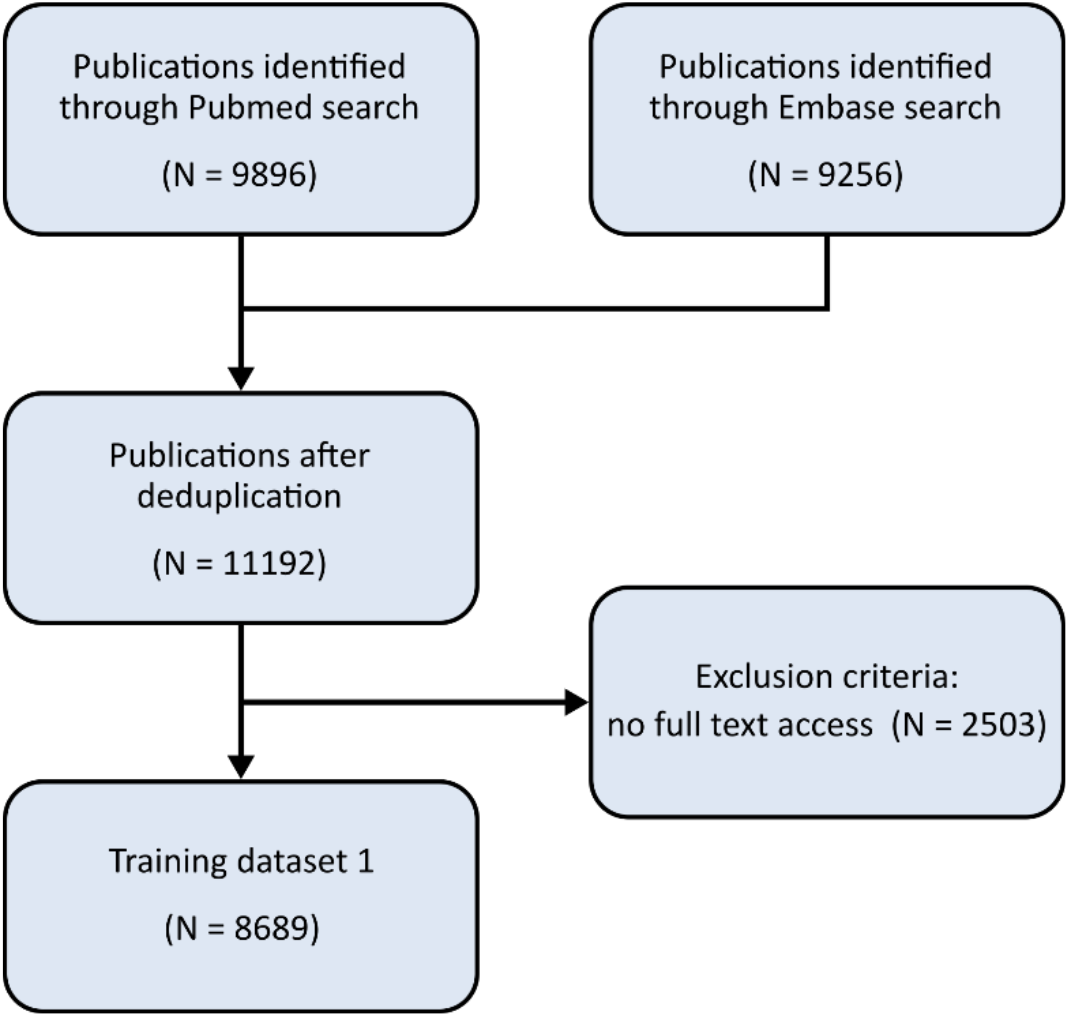
Inclusion and exclusion flowchart for training dataset 1.

The second training dataset consisted of publications randomly selected from all publications listed in PubMed in the year 2018. For this, we downloaded a list of all PMIDs from 2018 by exporting the search results using the following search filter:

- Publication date in 2018
- Language: English
- Publication type: Journal Article
- Publication type NOT: Editorial, Comment, Letter to Editor, Review, Erratum, Guideline, Historical Article

We retrieved a list of 1091518 PMIDs from which we randomly sampled 1500 (0.14%) IDs. We again downloaded the full texts and were able to retrieve 992 (66%) publications. As the PubMed metadata on the publication type was not always correct, we manually checked the publication types of the downloaded full texts. We excluded publications with the article types Editorial, Comment, Letter to Editor, Review, Erratum, Guideline, Historical Article, Case report. Additionally, we excluded other non-research articles that did not fit in any of those categories (e.g. Question & Answer) and research articles that were out of the scope of the biomedical literature (social science & history of science). After excluding those publication types, we were left with 868 publications (see Figure 2).

**Figure 2:**
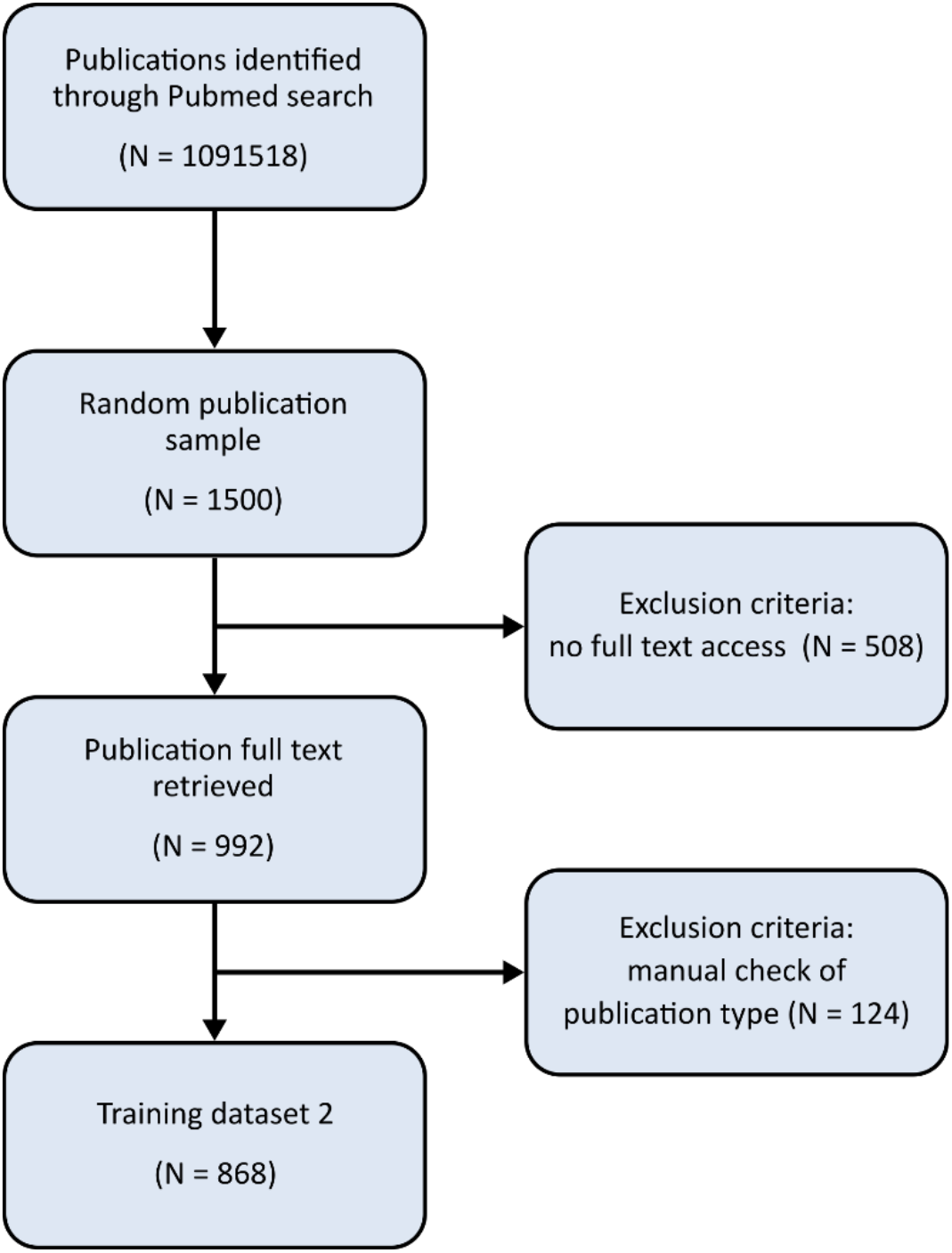
Inclusion and exclusion flowchart for training dataset 2.

This dataset was originally planned as the validation dataset, but due to low sensitivity of the algorithm developed on the institutional publication record, we decided to use this second dataset to improve the algorithm further. We then drew another sample of N = 1400 publications (downloaded full texts: 964, 69%, after publication type check: 792) listed in PubMed in the year 2018 that was used as the final validation dataset (see Figure 3).

**Figure 3:**
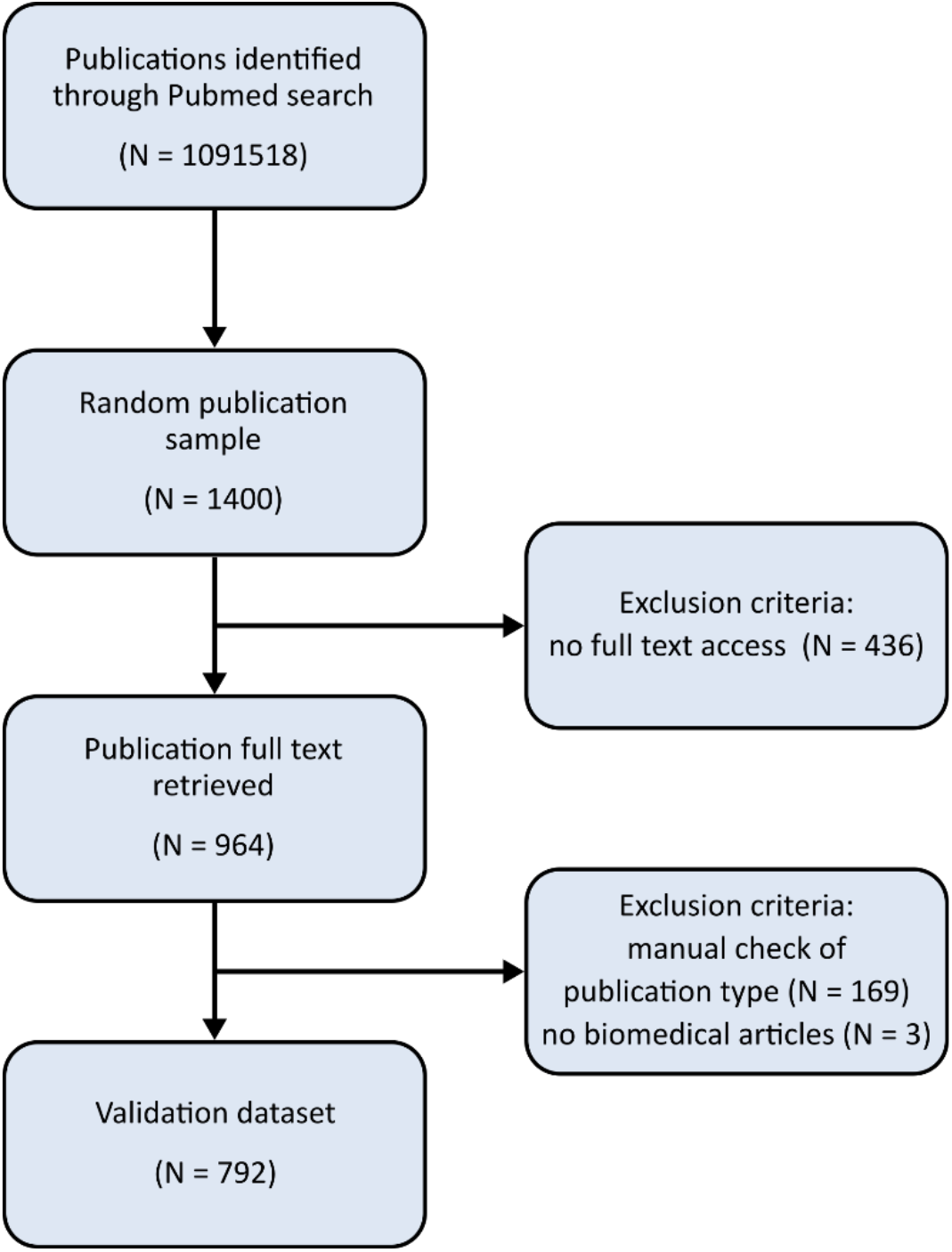
Inclusion and exclusion flowchart for the validation dataset.

### Data preprocessing

To prepare the publications for text mining we performed a number of preprocessing steps. Using an R script, we first converted the PDFs to text files using the poppler library (*Poppler* (version 0.67.0) 2018). We then loaded the text files into R and split the texts into the individual sentences using the “tokenizers” R package (Mullen et al. 2018). As the last step, the text was converted to lowercase.

### Manual check of test & validation samples

To establish a gold standard for the Open Data detection in the training and validation samples, we manually screened the publications for Open Data. The screening followed a standardized procedure, involving the following steps:

1. We checked if the article is a research article. Editorials, reviews, case reports or other article types (see above) were excluded from the sample. As the automatic exclusion via the metadata did not detect all those cases, this manual check was necessary.
2. We screened the article for a distinct “data availability” section, which is primarily found either at the beginning of the publication or at the end of the main text, before the references.
3. We used the search function to search for several keywords: “availab”, “access”, “raw”, “deposit”, “supp”, “data”, “code”, “script”, “software”. The sentences of the detected keywords were then manually checked for relevant Open Data or Open Code statements. If some of the keywords appeared more than ten times in the manuscript, they were replaced by more specific keywords: “access” was replaced by “accession” and “accessible”, “supp” was replaced by “supporting” and “supplementary”, and “data” was replaced by “dataset”, “data set”, “database” and” data base”. We used this approach instead of reading the entire full text for time reasons but also to find the most relevant passages more reliably.
4. If a relevant statement was found, we manually verified that the data or analysis code was indeed present and that they fulfilled the Open Data definition as explained above. If the found data were stored in a very large ZIP file (>200MB), we did not extract the data but assumed that they contained data that agree with the Open Data definition.

A more detailed description of the manual search strategy can be found in supplemental file S2.

To estimate the interrater reliability of the manual screening, a second rater checked 100 publications from the validation dataset independently. For this, we chose a random sample of 25% of all positively identified Open Data papers (n=21) and added a random sample of n=79 negatively identified papers to partially even out the strong class imbalance in this dataset. Both raters used the manual screening procedure as defined above to determine the Open Data status (either “true” or “false”) for each of the 100 publications independently. We then calculated the percentage of cases where the raters agreed with their assessment of the Open Data status of the paper. For details on the interrater sample see supplemental file S6.

### Algorithm development procedure

For the development of ODDPub, we used an iterative procedure to manually derive and refine a set of keywords and key phrases to detect Open Data statements, focusing on first increasing the sensitivity and subsequently increasing the specificity in each step. As we could not check all publications in our first training sample manually for Open Data and as the prevalence of Open Data was rather low, we focused on analyzing all positively detected cases as well as a random subset of the negatively detected cases during the first iterations. We give a brief summary of the development procedure here. A more detailed description can be found in supplemental file S3. For details on the different algorithm stages, see supplemental files S4 and S5 as well as the GitHub repository (https://doi.org/10.5281/zenodo.3760970).

We started with a small number of confirmed Open Data publications. We extracted all keywords or key phrases relevant for the Open Data status and wrote an R script that uses regular expression matching to search for the keywords in the sentences of the preprocessed publication texts. If any of the keywords was detected, the publication was counted as hit.

We applied this first version of the keyword search to the whole Charité training dataset (N=8689 publications). We manually checked samples from the positively detected cases as well as from the negative cases. Using the information from the false positive and false negative cases, we improved the algorithm by adding new keywords and forming keyword groups of which several have to match in the same sentence. This was repeated for three rounds, where the algorithm was again applied to the whole training dataset after each improvement and samples of positive and negative cases were manually assessed to further improve the keyword groups and to decrease the false positive and false negative rate of the algorithm.

We then applied the improved algorithm to the first random sample of 992 PubMed publications. First, we manually searched all publications in the sample for Open Data, where we detected 98 (9.9%) Open Data publications. Then we ran the algorithm on the same publication sample and compared the results to the gold standard of manual search. We found a sensitivity of 45% and a specificity of 98%. As we considered a sensitivity of 45% insufficient, we decided to use this first PubMed sample to improve the algorithm further and to draw a second PubMed sample as the final validation sample. In a final round of improvement, we used the manually detected Open Data statements from the first

PubMed sample that were missed by the algorithm to add new keywords and keyword categories for improved coverage of the algorithm. Additionally, we used the false positive cases to remove keywords that primarily led to false positive detections. Detailed information on the final version of ODDPub can be found in supplemental file S4.

### Measuring performance of the algorithm

We applied the final version of our algorithm to all publications of the validation dataset and compared the results with the manual screening results using the sensitivity, specificity and F1 score as performance measures.

As an additional benchmark, we also extracted the associated metadata on the datasets recorded in the databases PubMed and Web of Science for all publications in the validation sample. For PubMed we used the metadata field ‘Secondary Source ID’, which contains database names and dataset identifiers of datasets associated with the publications. Those databases comprise many genetic databases as well as the general-purpose databases Dryad and Figshare. For Web of Science we used the associated datasets that were linked to the publications of the validation sample. Those can be found as an additional linked field “Associated Data” under the title of a publication on the Web of Science website.

We calculated the number of datasets listed in the metadata field that were also found by ODDPub as well as the number of listed datasets that were missed by ODDPub.

## Results

### Structure of the resulting algorithm

The resulting text-mining algorithm has a hierarchical keyword structure and searches for several categories of similar keywords in each sentence (see Figure 4). We grouped similar keywords identified in the statements indicating open data or code from the training dataset into categories (see supplemental file S4). Examples are the categories FIELD_SPECIFIC_DB, containing database names like “Gene Expression Omnibus”, “Sequence Read Archive”, or “Protein Data Bank” and AVAILABLE, containing words to denote that the data have been made available, like “deposited”, “released”, or “submitted”. The explanations in this section hold true for the detection of both Open Data and Open Code.

Groups of several keyword categories were then formed that each comprise several keyword categories. Those groups of keyword categories cover different ways of data sharing like field-specific repositories, general-purpose repositories or deposition of data in the supplement (see Table 1). For a match in a group of keyword categories, at least one of the keywords in each underlying keyword category has to match. For the example of the “Field-specific databases” category group, we need matches for any of the keywords of the underlying categories FIELD_SPECIFIC_DB, AVAILABLE and ACCESSION_NR (which contains regular expressions to match the accession numbers of those databases). There are two types of keyword category groups. Either all keywords are searched within the same sentence or the keywords are searched across sentence borders, allowing for a maximum number of other words between the keywords. This is necessary in cases like “*S2 Table. Raw data. (XLS)*”, where the information is spread across sentence borders. The whole text of the publication is checked for a match in any of the keyword category groups. A publication is categorized as Open Data if any of the keyword category groups match for at least one passage in the full text. All details on the categories see supplemental file S4 or the GitHub repository (https://doi.org/10.5281/zenodo.3760970).

**Figure 4:**
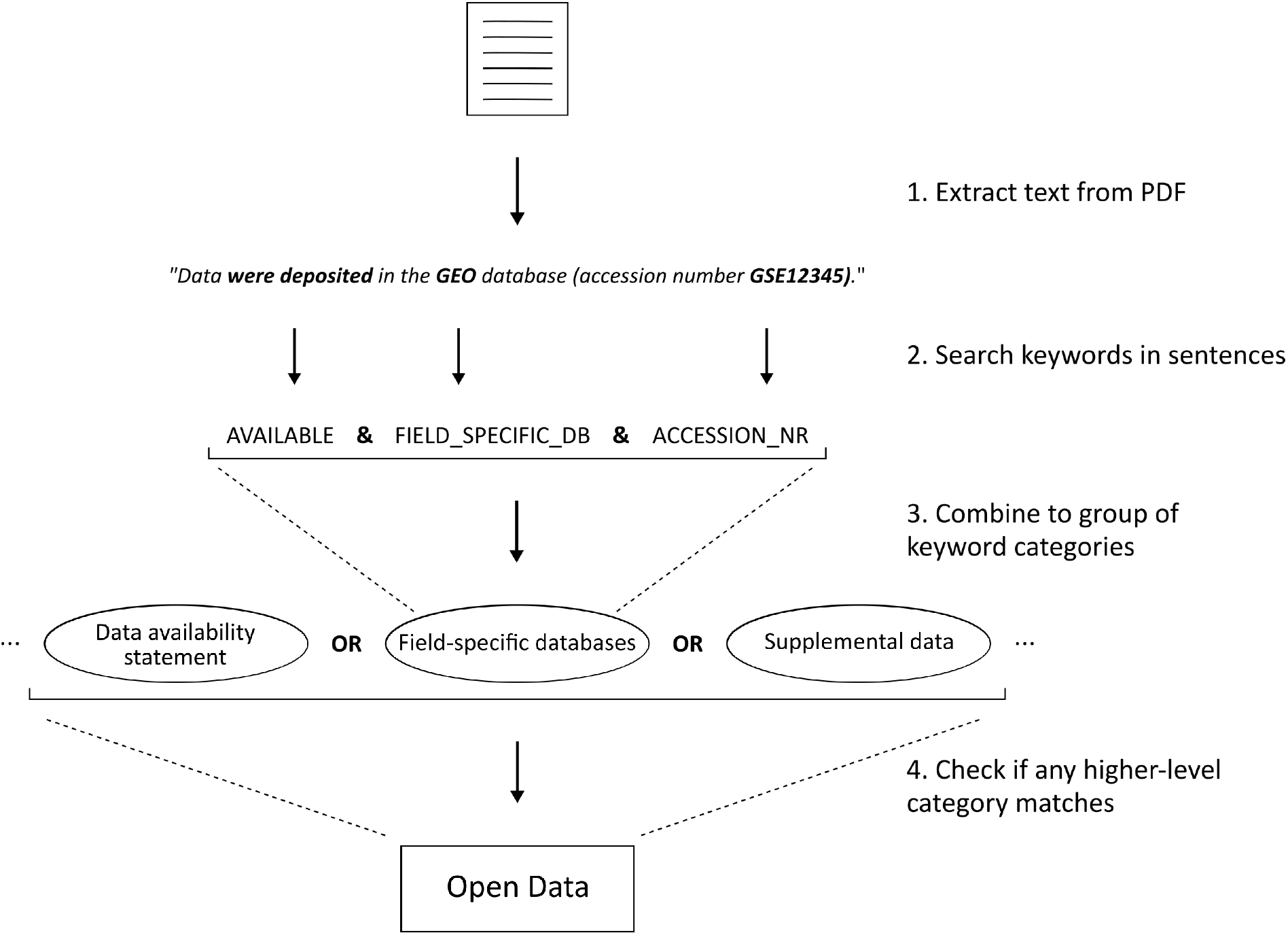
Schema of the algorithm workflow and architecture.

**Figure 5:**
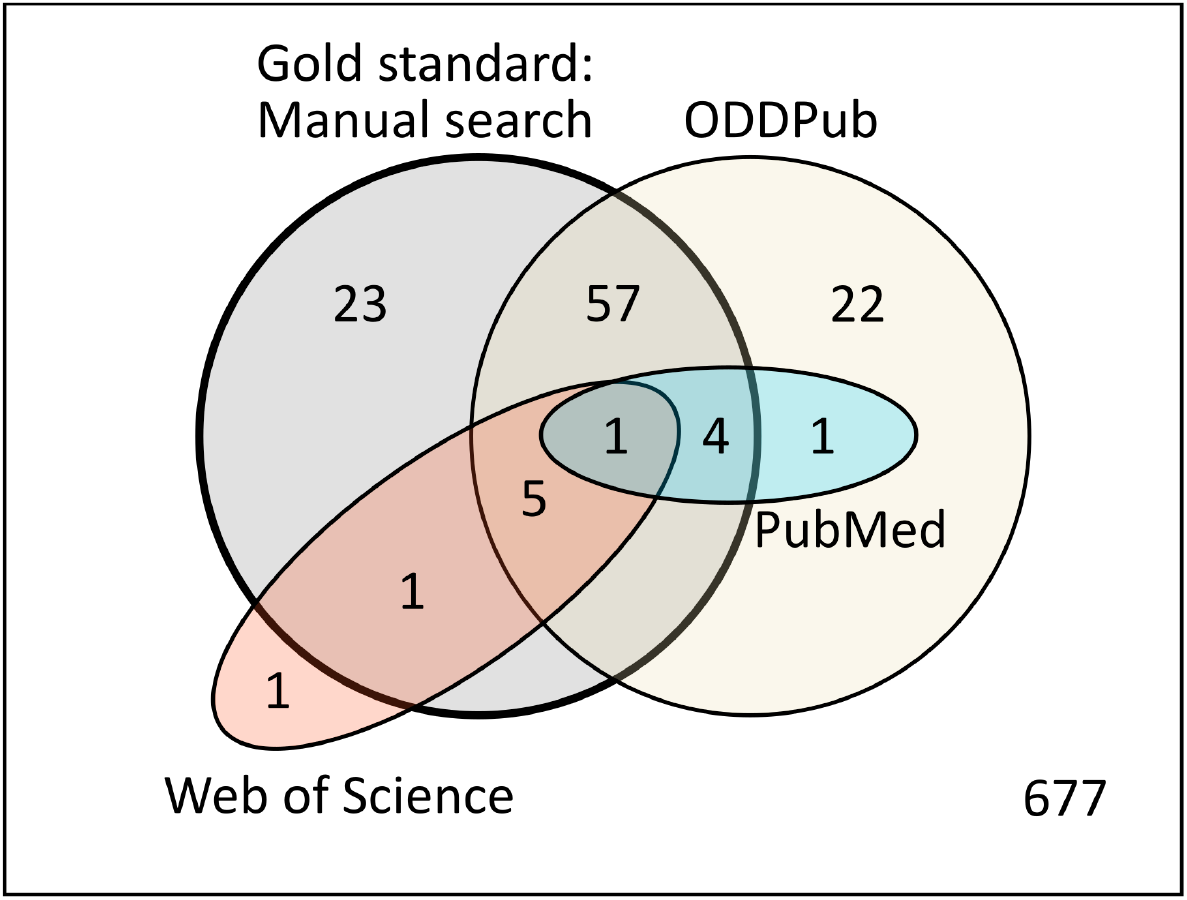
Venn diagram of the overlap between the detected Open Data publications for the four different detection methods on the validation dataset: Manual search, ODDPub, PubMed, and Web of Science. The manual search represents the gold standard. All 23 publications detected by ODDPub but not the manual search are false positive detections.

**Table 1:** Different keyword categories used by ODDPub to detect Open Data and Open Code.

| Combined Keyword Categories | Explanation |
| --- | --- |
| **Open Data** |  |
| Field-specific repositories | Checks if data deposition in field-specific database and accession number is mentioned |
| General-purpose repositories | Checks if data deposition in general-purpose database that uses no accession number is mentioned |
| Dataset | Checks if dataset that is given specific number (e.g. “dataset S1”) is mentioned |
| Supplemental table or data | Checks if a numbered file or table or raw data is mentioned together with specific file formats |
| Supplementary raw/full data with specific file format | Checks if raw/full data is mentioned together with specific file formats |
| Data availability statement | Checks if an accession number or a repository name is mentioned in the data availability section |
| Dataset on GitHub | Checks if data deposition on GitHub is mentioned |
| Data journals | Checks journal DOI for certain known data journals |
| **Open Code** |  |
| Source-code availability | Checks if availability of source code is mentioned |
| Supplementary Source-code | Checks if source code in the supplement is mentioned |

In addition to the Open Data status, the algorithm outputs the sentence in which the Open Data statement was detected. This allows a fast manual check of the results in cases where removal of the remaining false positives is crucial. Also, it allows to use the detected Open Data statements for further analysis (e.g. on details of how data are shared).

### Performance criteria

In this section, we report the performance measures of ODDPub for the validation dataset. For this, we compared the results from the manual screening to the results obtained with ODDPub.

In the manual search Open Data was detected for 11.5% (n=91) of the publications in the validation dataset and Open Code was detected for 1.4% (n=11) of the publications.

The results of the Open Data prediction for ODDPub compared to the gold standard of manual screening can be found in Table 2. We obtained a sensitivity of 0.73, a specificity of 0.97, and a F1-score of 0.73. The results of the Open Code prediction for ODDPub compared to the gold standard of manual screening yield a sensitivity of 0.73, a specificity of 1.00, and a F1-score of 0.64 (see Table 3).

**Table 2:** Predictions of ODDPub for Open Data on the validation dataset in comparison to the manual screening.

| Open Data |  | ODDPub |  |
| --- | --- | --- | --- |
|  |  | Yes | No |
| Human rater | Yes | 67 | 24 |
|  | No | 23 | 678 |

**Table 3:** Predictions of ODDPub for Open Code on the validation dataset in comparison to the manual screening.

| Open Code |  | ODDPub |  |
| --- | --- | --- | --- |
|  |  | Yes | No |
| Human rater | Yes | 8 | 3 |
|  | No | 6 | 775 |

To check if any Open Data publication found by ODDPub was missed by the manual screening, we manually checked the Open Data statements of all 24 publications detected with ODDPub but not with the manual screening. We identified one of those 24 publications as Open Data publication according to our definition and counted it as an Open Data publication in Table 2. The remaining 23 publications turned out to be false positive detections (see Table 5).

As only few cases of Open Code were found, the estimates for the performance metrics for the Open Code detection are unreliable. Two Open Code publications were only found by ODDPub and not by the manual screening. As for the case of Open Data, we manually verified the Open Code status of these publications and counted them as Open Code publications in Table 3. Of the six false positive cases, there are five cases of Open Code reuse.

### Comparison to PubMed and Web of Science

Next, we compared our results to the linked datasets found in the databases PubMed and Web of Science.

For the comparison with PubMed we retrieved the entries in the ‘ Secondary Source ID’ field in PubMed for the 792 publications in the validation sample. We found that 19 of those publications have an entry in the secondary ID field. 11 of the entries referred to clinical trial registration IDs (8 on ClinicalTrials.gov, 3 on ISRCTN). Four entries linked to a Figshare dataset, two linked to a Dryad repository, and one entry linked to the Genbank and the Protein Data Bank, respectively. All Figshare and Dryad datasets were also detected by ODDPub. The two entries for Genbank and the Protein Data Bank were both cases of reuse of previously published data. Those do not fit our definition of Open Data and were not detected by ODDPub.

For the comparison with Web of Science, we retrieved the datasets linked in Web of Science for the 792 publications in the validation samples. We found that 11 publications have reported associated data. Those fall into the following categories:

- 3 of them were previously published R packages on CRAN (no dataset; also did not fall under our definition of Open Code)
- 4 had links to datasets in the ‘ Cambridge Crystallographic Data Centre’ (CCDC) database
- 2 had links to datasets in the ‘ Protein Data Bank’ (PDB) database
- 1 had a link to a Dryad repository
- 1 had a link to a dataset in the PRIDE database

Of those 8 reported datasets, 6 were found by both ODDPub and manual search. The PDB dataset was only found by manual search and one of the CCDC datasets was found by neither ODDPub nor the manual search. Only the link to the Dryad repository could be found through both PubMed and Web of Science.

In summary, both PubMed and Web of Science linked only few of the datasets found by ODDPub for the publications in the validation sample.

### Interrater reliability

For the 100 publications of the validation dataset that were checked by two independent raters, an agreement regarding Open Data status was reached in 95% of the cases (see supplemental Table S6).

### Categories of Open Data detected

To determine which categories of Open Data were most frequently discovered, we categorized all 91 manually detected Open Data publications of the validation dataset in the following categories: supplemental data, field-specific repository, general-purpose repository (including GitHub), institutional repository, data journal, and personal/project-specific website. One publication can be assigned to multiple categories if applicable.

Most of the detected Open Data publications used either the supplement or a field-specific repository (Table 4). Of the 24 manually detected Open Data cases that were not found by ODDPub, there were 17 cases with supplemental data and 7 cases with field-specific repositories. Sharing data via the supplement was thus less likely to be detected by the algorithm.

**Table 4:** Types of data sharing observed in the manually detected Open Data publications of the validation sample.

| Category | Number of occurrences |
| --- | --- |
| Supplemental Data | 42 |
| Field-specific repository | 40 |
| General-purpose repository (including GitHub) | 14 |
| Institutional repository | 0 |
| Personal/project-specific website | 1 |
| Data journal | 0 |

Additionally, we categorized the 23 false positive cases detected by ODDPub in the validation sample (see Table 5). Most of the cases were not labeled as Open Data in the manual search because our definition of Open Data was not fulfilled in some way. In three cases, the algorithm made a major mistake and detected a statement that was not related to data sharing at all. In one case, an Open Data publication that was missed by the manual search was detected.

**Table 5:**
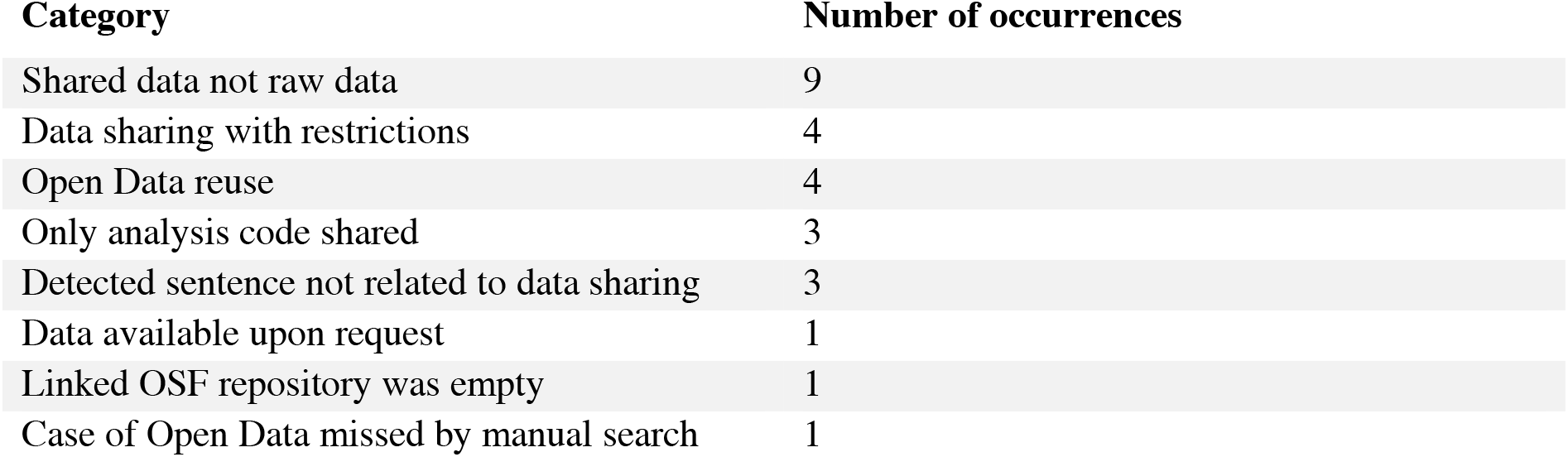
Reasons for false positive cases detected by ODDPub in the validation sample.

A list of all detected Open Data statements for the training and validation datasets as well as the results of the manual screening can be found in a separate OSF repository (https://doi.org/10.17605/OSF.IO/YV5RX).

## Discussion

In this work, we developed a text-mining algorithm that can screen biomedical publications for mentions of Open Data sharing, as well as the provision of self-developed code (Open Code). Our text mining algorithm allows to screen large numbers of publications for statements indicating Open Data sharing automatically.

An advantage of our approach compared to other studies that focus on specific journals that have a dedicated data availability statement (Federer et al. 2018; Colavizza et al. 2020) is the broader applicability. As our algorithm screens all parts of a publication, Open Data statements can be detected anywhere in the text. This allows screening of publications in journals without dedicated data availability statements. A disadvantage of this approach is that this makes the task more complex compared to an approach that only focuses on the data availability statements. The studies of (Federer et al. 2018; Colavizza et al. 2020) both could use a less complex classification algorithm on the data availability statements, which is not possible when searching the full text.

Compared to the two currently available publication databases PubMed and Web of Science that do link information on Open Data to publications we could show a much better coverage of our algorithm on the validation dataset. This suggests that currently our approach is superior to the other available approaches for the automated detection of Open Data publications from a larger publication corpus.

### Applications

ODDPub has several possible areas of application. One area is the large-scale assessment of data sharing rates, for example on the level of different subject areas, where a manual assessment is too time-consuming and where a strategy relying on data availability statements alone is not possible due to many journals that have not yet implemented those statements.

Another possible area is the detection of individual Open Data publications in a larger publication corpus, e.g. to identify the Open Data publications of a specific research institution. This can be necessary if the institution wants to distribute additional performance oriented funding for publications with Open Data. In this case, the output of the detected Open Data statements by the algorithm helps to gain more details on the detected publications and allows for a fast manual check for false positives. We used ODDPub in a semi-automated way (automated screening plus manual check) for the detection of Open Data publications in the Charité publication body among researchers eligible for the annual allocation of performance-based funding. In 2019 over 120.000 Euros were allocated among Charité researchers based on semi-automated Open Data detection, and this is planned to be continued in 2020 (Kip et al. 2019; BIH 2019).

### Limitations

Open Data is a fast-evolving field, with some but yet few standards. Therefore, the provided results on specificity and sensitivity are a snapshot in time, and might change quickly. The algorithm will require further and continuous development and adaption to the changes and developments in the field, e.g. new repositories and the wider adoption of data availability statements.

One of the main limitations is the sensitivity of 73% obtained for the current version of ODDPub. The missed Open Data publications might be problematic in applications where a very high coverage of Open Data publications is important. As most of the missed Open Data publications shared supplemental data, this problem might be reduced in the future if more research data are shared through repositories.

Our approach focuses on the detection of open research data that are linked to a publication. For an assessment of datasets published independently of a publication, a different approach would be necessary.

Additionally, the scope of our algorithm was the biomedical literature and many of the Open Data statements found by the algorithm refer to repositories specific to the biomedical domain. The performance on other research areas has not been tested and one can expect a decrease in performance of the algorithm when applied to other research areas. The accessibility of journals is another limitation. As not all journals were accessible to us, we could not include those into our training dataset. Due to this, we could have potentially introduced a bias in our training dataset and missed additional relevant repositories or keywords.

Importantly, our definition of Open Data did not require data to be FAIR (Wilkinson et al. 2016) or complete. Thus, even in the manual check, which we used as a gold standard, the standards of data provision were set very low. Indeed, while we could not quantify the fraction of reported findings for which raw data had been shared, providing the data underlying all results was surely rather exception than norm. Similarly, standards of machine-readability and documentation in our Open Data sample clearly fell short of multiple FAIR criteria for most datasets, even though this was not systematically assessed. Thus, Open Data sharing rates reported here will not be comparable with those detected applying more stringent criteria of completeness and reusability, which would be recommendable to do as Open Data sharing becomes more common and standardized.

For the detection of Open Code, one obstacle is that the standardization of Open Code statements is still lower than for Open Data (e.g. no database IDs). Both the lower standardization and the lower prevalence of Open Code combined make it difficult to develop a robust tool for Open Code statement detection.

### Outlook

How Open Data is deposited and linked will change over time, with new data repositories emerging and changing file formats. Thus, the algorithm needs to be adapted in the future to keep up with these developments. One possibility would be to update all repositories listed in e.g. re3data.org and FAIRsharing.org automatically to keep the algorithm up to date regarding available repositories. However, ODDPub was developed given the current landscape of Open Data, in which solutions for reusable Open Data sharing and standardized, automated detection of the datasets are already present, but have not yet been universally adopted. These include in particular

i. sharing of datasets through repositories rather than as supplementary materials, leading to both easier findability and citation as independent research outputs
ii. adoption of the Joint Declaration of Data Citation Principles (Martone 2014), leading to datasets being cited in the reference list
iii. the inclusion of data availability statements in journal articles, which can clarify, which of the cited datasets are primary data, created or collected by the authors

Another relevant point concerning the citation of datasets in the future is the distinction between citations of newly generated datasets and of re-used datasets. In our screening, we observed that Open Data reuse can create confusion with the sharing of primary data when detecting these practices by text mining. Thus, data citation principles and corresponding metadata fields in registries need to assure that the sharing of primary data and the reuse of data can be distinguished. In addition, granting access to data under restrictions is another important way of data sharing especially in biomedical research. Practices of definition, annotation and detection need to consider this, and provide the necessary information in a machine-readable fashion, if the current definition of Open Data is to be maintained.

In conclusion, the text-mining algorithm ODDPub presented in this study can detect sharing of raw data (‘Open Data’) in publications from the biomedical literature. It can detect different ways of sharing Open Data, like field-specific repositories or sharing of raw data in the supplement. The results of the validation of the algorithm indicate that it finds the majority of Open Data publications in the current system of how Open Data are shared and linked in the biomedical literature. In the future, our method can be used to assess data sharing rates in the biomedical literature on a larger scale, which would not be possible with a manual approach.

## Supporting information

Supplement S1: Search queries to retrieve the publications

Supplement S2: SOPs for manual screening of publications for Open Data

Supplement S3: Detailed description of the algorithm development procedure

Supplement S4: ODDPub Version History

Supplement S5: Detailed evaluation of the results of each algorithm development stage

Supplement S6: Interrater sample

## Acknowledgements

We thank Meggie Danziger, Maxim Benz, Claudia Kirchert and especially Yasmin Aktas for their help with the manual screening of publications. Additionally, we thank Lisa Liebenau, Florian Beng and Clemens Blümel for the helpful discussions in the early conceptual phase of this project. Finally, we thank Benjamin Gregory Carlisle and Ulrich Dirnagl for their helpful comments on our manuscript.

## Data Accessibility Statement

The ODDPub algorithm is available as R package on GitHub (https://doi.org/10.5281/zenodo.3760970, RRID:SCR_018385) under the MIT license. A dataset containing information on the publications used in the training and validation datasets, including the publication DOIs, the results of the manual screening and the extracted Open Data statements can be found under https://doi.org/10.17605/OSF.IO/YV5RX.

## Supplemental Material

**S1 –** Search queries for the databases PubMed and Embase.

**S2 –** Detailed description of the procedure for the manual search for Open Data statements in the publications.

**S3 –** Detailed description of the algorithm development procedure.

**S4 –** Table with a detailed description of the keywords used in each of the different versions of ODDPub.

**S5 –** Table with the detailed evaluation of the results of each algorithm development stage that were used to inform the next version of the algorithm.

**S6 –** Results of the interrater reliability sample.

## Bibliography

BIH. 2019. “BIH Rewards Open Data in an Effort to Make Science More Verifiable.” June 20, 2019. https://www.bihealth.org/en/notices/bih-rewards-open-data-in-an-effort-to-make-science-more-verifiable/.

Colavizza, Giovanni, Iain Hrynaszkiewicz, Isla Staden, Kirstie Whitaker, and Barbara McGillivray. 2020. “The Citation Advantage of Linking Publications to Research Data.” ArXiv:1907.02565 [Cs], March. http://arxiv.org/abs/1907.02565.

Cousijn, Helena, Amye Kenall, Emma Ganley, Melissa Harrison, David Kernohan, Thomas Lemberger, Fiona Murphy, et al. 2018. “A Data Citation Roadmap for Scientific Publishers.” Scientific Data 5 (1): 180259. https://doi.org/10.1038/sdata.2018.259.

“EOSC Declaration.” 2017. https://ec.europa.eu/research/openscience/pdf/eosc_declaration.pdf.

EUA. 2017. “Towards Open Access to Research Data.” https://eua.eu/downloads/publications/towards%20open%20access%20to%20research%20data%20aims%20and%20recommendations%20for%20university%20leaders.pdf.

Fecher, Benedikt, Sascha Friesike, and Marcel Hebing. 2015. “What Drives Academic Data Sharing?” PLOS ONE 10 (2): e0118053. https://doi.org/10.1371/journal.pone.0118053.

Federer, Lisa M., Christopher W. Belter, Douglas J. Joubert, Alicia Livinski, Ya-Ling Lu, Lissa N. Snyders, and Holly Thompson. 2018. “Data Sharing in PLOS ONE: An Analysis of Data Availability Statements.” PLOS ONE 13 (5): e0194768. https://doi.org/10.1371/journal.pone.0194768.

Guedj, D, and C Ramjoué. 2015. “European Commission Policy on Open-Access to Scientific Publications and Research Data in Horizon 2020.” Biomedical Data Journal 01 (1): 11–14. https://doi.org/10.11610/bmdj.01102.

Kip, Miriam, Evgeny Bobrov, Nico Riedel, Heike Scheithauer, Thomas Gazlig, and Ulrich Dirnagl. 2019. “Einführung von Open Data Als Zusätzlicher Indikator Für Die Leistungsorientierte Mittelvergabe (LOM) Forschung an Der Charité – Universitätsmedizin Berlin,” October. https://doi.org/10.5281/ZENODO.3511191.

Martone, M, ed. 2014. “Data Citation Synthesis Group: Joint Declaration of Data Citation Principles.” San Diego CA: FORCE11. https://doi.org/10.25490/a97f-egyk.

Milham, Michael P., R. Cameron Craddock, Jake J. Son, Michael Fleischmann, Jon Clucas, Helen Xu, Bonhwang Koo, et al. 2018. “Assessment of the Impact of Shared Brain Imaging Data on the Scientific Literature.” Nature Communications 9 (1): 2818. https://doi.org/10.1038/s41467-018-04976-1.

Mullen, Lincoln A., Kenneth Benoit, Os Keyes, Dmitry Selivanov, and Jeffrey Arnold. 2018. “Fast, Consistent Tokenization of Natural Language Text.” Journal of Open Source Software 3 (23): 655. https://doi.org/10.21105/joss.00655.

Naudet, Florian, Charlotte Sakarovitch, Perrine Janiaud, Ioana Cristea, Daniele Fanelli, David Moher, and John P A Ioannidis. 2018. “Data Sharing and Reanalysis of Randomized Controlled Trials in Leading Biomedical Journals with a Full Data Sharing Policy: Survey of Studies Published in *The BMJ* and *PLOS Medicine*.” BMJ, February, k400. https://doi.org/10.1136/bmj.k400.

NIH. 2003. “NIH Data Sharing Policy and Implementation Guidance.” https://grants.nih.gov/grants/policy/data_sharing/data_sharing_guidance.htm.

Open Knowledge Foundation. 2017. “Open Definition 2.1 - Open Definition - Defining Open in Open Data, Open Content and Open Knowledge.” https://opendefinition.org/od/2.1/en/.

Piwowar, Heather A., and Todd J. Vision. 2013. “Data Reuse and the Open Data Citation Advantage.” PeerJ 1 (October): e175. https://doi.org/10.7717/peerj.175.

Poppler (version 0.67.0). 2018. https://poppler.freedesktop.org/.

Pronk, Tessa E. 2019. “The Time Efficiency Gain in Sharing and Reuse of Research Data.” Data Science Journal 18 (March): 10. https://doi.org/10.5334/dsj-2019-010.

Rowhani-Farid, Anisa, and Adrian G Barnett. 2016. “Has Open Data Arrived at the *British Medical Journal (BMJ)*? An Observational Study.” BMJ Open 6 (10): e011784. https://doi.org/10.1136/bmjopen-2016-011784.

Savage, Caroline J., and Andrew J. Vickers. 2009. “Empirical Study of Data Sharing by Authors Publishing in PLoS Journals.” PLoS ONE 4 (9): e7078. https://doi.org/10.1371/journal.pone.0007078.

“Sorbonne Declaration on Research Data Rights.” 2020.

Stodden, Victoria, Peixuan Guo, and Zhaokun Ma. 2013. “Toward Reproducible Computational Research: An Empirical Analysis of Data and Code Policy Adoption by Journals.” PLoS ONE 8 (6): e67111. https://doi.org/10.1371/journal.pone.0067111.

Strech, Daniel, Tracey Weissgerber, Ulrich Dirnagl, and on behalf of QUEST Group. 2020. “Improving the Trustworthiness, Usefulness, and Ethics of Biomedical Research through an Innovative and Comprehensive Institutional Initiative.” PLOS Biology 18 (2): e3000576. https://doi.org/10.1371/journal.pbio.3000576.

“The Berlin Declaration on Open Access to Knowledge.” 2003. https://openaccess.mpg.de/Berliner-Erklaerung.

The Concordat Working Group. 2016. “Concordat on Open Research Data.” https://www.ukri.org/files/legacy/documents/concordatonopenresearchdata-pdf/.

Tudur Smith, C, C Hopkins, M Sydes, K Woolfall, M Clarke, G Murray, and P Wiliamson. 2015. “Good Practice Principles for Sharing Individual Participant Data from Publicly Funded Clinical Trials.” https://www.methodologyhubs.mrc.ac.uk/files/7114/3682/3831/Datasharingguidance2015.pdf.

Vines, Timothy H., Arianne Y.K. Albert, Rose L. Andrew, Florence Débarre, Dan G. Bock, Michelle T. Franklin, Kimberly J. Gilbert, Jean-Sébastien Moore, Sébastien Renaut, and Diana J. Rennison. 2014. “The Availability of Research Data Declines Rapidly with Article Age.” Current Biology 24 (1): 94–97. https://doi.org/10.1016/j.cub.2013.11.014.

Wilkinson, Mark D., Michel Dumontier, IJsbrand Jan Aalbersberg, Gabrielle Appleton, Myles Axton, Arie Baak, Niklas Blomberg, et al. 2016. “The FAIR Guiding Principles for Scientific Data Management and Stewardship.” Scientific Data 3 (1): 160018. https://doi.org/10.1038/sdata.2016.18.

