## Supplement S1: Search queries to retrieve the publications for "ODDPub – a Text-Mining Algorithm to Detect Data Sharing in Biomedical Publications"

**PubMed search:**

We used the following search query for PubMed

*(charit*[Affiliation] OR
university medicine berlin[Affiliation] OR
universitatsmedizin berlin[Affiliation] OR
(campus[Affiliation] AND virchow[Affiliation] AND berlin[Affiliation]) OR
(campus[Affiliation] AND franklin[Affiliation] AND berlin[Affiliation]) OR
(campus[Affiliation] AND mitte[Affiliation] AND berlin[Affiliation])
) AND "2015/01/01"[Date - Publication]: "2017/12/31"[Date – Publication]*

As the “charit*” search string is too broad, we went through the search details that are shown on the Pubmed website after entering the search and selected all search terms from the list that were used by PubMed for the wildcard search for “charit*” that we do not consider related with the Charité. Publications containing the following strings in the affiliation are filtered out in the subsequent step:

"charit6", "charita", "charita74", "charitable", "charitac", "charitakis", "charitakisc", "charitansky", "charitas", "charitat", "charith", "charitha", "charithalg", "charithhanduwalage", "chariti", "charitidi", "charitidou", "charitie", "charities", "charitiota", "charitle", "charitn", "charito54", "charitojm", "chariton", "charitonia", "charitonidi", "charitos", "charitt", "charitte", "charitx00e9", "charity", "charity'", "charity''", "charity's", "charityann80", "charitybomi", "charityhungu", "charityjadeknight", "charityjuang", "charitymemorialclinic", "charityzxh", "charitz"

**Embase search:**

We used the following search query for Embase

(charit* OR university medicine berlin OR universitatsmedizin berlin OR (campus AND virchow AND berlin) OR (campus AND franklin AND berlin) OR (campus AND mitte AND berlin)).in AND (2015 OR 2016 OR 2017).yr
