## Supplement S2: SOPs for manual screening of publications for Open Data for "ODDPub – a Text-Mining Algorithm to Detect Data Sharing in Biomedical Publications"

Supplement S2: SOPs for detecting publications with Open Data

### 1. Manual Screening for Open Data

These SOPs (Standard Operating Procedures) were developed to capture the procedure for detecting Open Data. Specifically, the aim is to identify publications that openly shared their research data.

#### Procedure

1. If the HTML version of the publication is screened instead of the PDF version: make sure that not only the abstract but also the full publication is displayed.
2. Check if the article is indeed a research article – there are some cases of editorials, reviews or other types of articles in the sample. In addition, case reports are not considered as research article in this context, as there are typically no research data that could be shared.
3. Check if there is a section “data availability”. For this, the first page as well as the end of the manuscript (before the references) are screened, as these are the positions at which this section appears most often. Additionally, search for the keyword “availab”. If a data availability section was found, check it for an Open Data statement. If no reference to shared data was found there, continue with the next step.
4. Keyword search. Search for the following keywords and check all detected sentences for an Open Data statement.
   a. “data” OR if this keyword appears more than 10 times, restrict to search for
      i. “dataset”
      ii. “database”
   b. “access” OR if this keyword appears more than 10 times, restrict to search for
      i. “accession”
      ii. “accessible”
   c. “raw” (possibly restrict search to whole words only)
   d. “supp” OR if this keyword appears more than 10 times, restrict to search for
      i. “supporting”
      ii. “supplementary”
   e. “deposited”
   f. Search for keywords for Open Code: “code”, “script”, “software”
5. If a data sharing statement was found, check it regarding the following criteria:
   a. Clear reference to the data: In the publication, it must be clearly mentioned that raw data are shared. A general note that e.g. there are „additional information” in the supplement is not sufficient.
   b. Data shared not only for illustrative purposes: For images or audiovisual data: those should not have only an illustrative character.
   c. Data are separate from main article: The data have to be shared separate from the main journal article, (typically in a repository or as supplemental material). Data that are contained within the article as tables are not counted as Open Data, as they are not easily findable (no own DOI or document as in the case of supplemental materials) and not machine readable (as they are typically in PDF or HTML format).
   d. Data are findable: If the data are not directly associated with the article as part of the supplement, a requirement for the findability of the data is a persistent identifier. Examples for this are DOIs, URLs or a database accession number.
      i. We accept if the data are findable via the accession number using a search engine like Google, if no direct link is given in the publication
      ii. Only mentioning the repository in which the dataset is deposited without giving the accession number is not accepted as Open Data
      iii. A link to a personal website is also not sufficient unless it is directly apparent without further search where the dataset is located on the website
   e. Data allow for replication or new analysis: at least one of the two conditions must be fulfilled:
      i. Data allow for the analytical replication of at least parts of the results presented in the article
      ii. Data allow for new analyses. Here, it is especially difficult to draw a line, but as a point of reference, it should hold that the data allow answering scientifically relevant questions.
   f. Genetic studies: For genetic studies, lists of significant genes from a microarray analysis are not sufficient.
   g. Patient data: Tables with epidemiological information on individual patients like age, sex, type of disease, comorbidities are only sufficient if those are the information that were actually analyzed in the study. It is however not sufficient if the information were only used to characterize the cohort.
   h. Smaller files (<50MB)
      i. If the technical possibility exists to open at least some of the provided files: check if the data contained are sufficiently raw, e.g. if values are given for individual runs/animals/patients.
      ii. If it is not technically possible to open the file (e.g. for uncommon file formats): if we can consider the data as Open Data depends on the clarity of the description of the data in the articles and the professionalism of the repository. A general assessment is difficult in this case. But if the data are clearly labeled as “raw” or if they are contained in an established field-specific repository, the data should be considered as raw data.
   i. Larger files (>50MB): If only larger files are available, we generally assume that the data are raw enough, as long as the description in the publication suggests that the data are raw. However, we do not download and check the data in those cases.
   j. ZIP-Files: For ZIP files it is often useful to download them and examine the contents, unless the files are very large (see previous point).
