## Supplement S3: Detailed description of the algorithm development procedure for "ODDPub – a Text-Mining Algorithm to Detect Data Sharing in Biomedical Publications"

Supplement S3: Algorithm development procedure

For the development of ODDPub, we used an iterative procedure to manually derive and refine a set of keywords and key phrases to detect Open Data statements, focusing on first increasing the sensitivity and subsequently increasing the specificity in each step. As we could not check all publications in our first training sample manually for Open Data and as the prevalence of Open Data is rather low, we focused on analyzing all positively detected cases as well as a random subset of the negatively detected cases during the first iterations. For details on the different algorithm stages, see supplemental files S4 and S5 as well as the GitHub repository (https://doi.org/10.5281/zenodo.3741404).

We started with a set of seven publications that contained Open Data. Prior to this project, these seven publications received an Open Data Award given by the authors’ institution and hence had already been confirmed as Open Data publications. These seven publications were therefore chosen as a plausible and pragmatic starting point. We extracted all keywords or key phrases relevant for the Open Data status (for detailed keywords, see supplemental files S4 and S5). We wrote an R script that uses regular expression matching to search for the keywords in the sentences of the preprocessed publication texts. If any of the keywords was detected, the publication was counted as hit.

We applied this first version of the keyword search to the whole Charité training dataset (N=8689 publications) which led to 1279 positive detected cases. From those cases, two independent raters manually checked for a random subset of 10% (N=120) whether the statements indicated Open Data by the definition given above. We found that 27 publications (23%) did agree with the definition, while the remaining 93 publications were false positives.

Using the results from the first manually checked sample, a second set of keywords was developed. For this, all positively identified Open Data publications from the first sample were selected and all relevant sentences that contained information on Open Data availability were extracted from these publications. From all identified relevant sentences, combinations of 2 or 3 keywords that appeared in the same sentence and that were distinctive for Open Data availability were extracted. Subsequently, we grouped the extracted keywords thematically and identified typical combinations of different groups of keywords. The revised second version of the keyword search is then the combination of all identified keyword groups.

Testing the second version of the keyword search on the same 10% sample gives a high sensitivity (93%), but still detects a high rate of false positives (positive predictive value, PPV = 57%). In the third round of the keyword refinement, the focus was on the false positive publications that do not contain Open Data but were still labeled as Open Data. All sentences that were marked as Open Data by version 2 of the classification algorithm were extracted from these papers. Using these sentences, several common patterns that led to the wrong identification as Open Data were identified and removed from the keyword search in version 3.

For a final round of improvement on this first training dataset, we ran version 3 of the keyword search on the whole Charité training dataset (N=8689 publications). Open Data was detected for 469 publications. We manually checked all positively classified as well as a random sample of 260 negatively classified publications (excluding publications from the first random sample). Based on those results, keywords and keyword categories that led to many false positives were removed while keywords for missed Open Data statements were added to the algorithm (ODDPub Ver. 4).

As we detected a number of statements referring to open analysis code but not to Open Data, we decided to form a separate keyword category for the detection of open analysis code statements.

We then applied ODDPub version 4 to the first random sample of 992 PubMed publications. First, we manually searched all publications in the sample for Open Data, where we detected 98 (9.9%) Open Data publications. Then we ran ODDPub version 4 on the same publication sample and compared the results to the gold standard of manual search. We found a sensitivity of 45% and a specificity of 98%. As we considered a sensitivity of 45% insufficient, we decided to use this first PubMed sample to improve the algorithm further and to draw a second PubMed sample as the final validation sample. In a final round of improvement, we used the manually detected Open Data statements from the first PubMed sample that were missed by ODDPub version 4 to add new keywords and keyword categories for improved coverage of the algorithm. Additionally, we used the false positive cases to remove keywords that primarily led to false positive detections. Detailed information on the final version of ODDPub can be found in supplemental file S4.
